# Angular and Linear Speed Cells in the Parahippocampal Circuits

**DOI:** 10.1101/2021.01.28.428631

**Authors:** Davide Spalla, Alessandro Treves, Charlotte N. Boccara

**Affiliations:** Sissa, Via Bonomea 265, 34136 Trieste, Italy; University of Oslo, Sognsvannsveien 9 Domus Medica, 0372 Oslo, Norway

## Abstract

An essential role of the hippocampal region is to integrate information to compute and update representations. How this transpires is highly debated. Many theories hinge on the integration of self-motion signals and the existence of continuous attractor networks (CAN). CAN models hypothesise that neurons coding for navigational correlates – such as position and direction – receive inputs from cells conjunctively coding for position, direction and self-motion. As yet, such conjunctive coding had not been found in the hippocampal region. Here, we report neurons coding for angular and linear velocity, distributed across the medial entorhinal cortex, the presubiculum and the parasubiculum. These self-motion neurons often conjunctively encoded position and/or direction, yet lacked a structured organisation, calling for the revision of current CAN models. These results offer insights as to how linear/angular speed – derivative in time of position/direction – may allow the updating of spatial representations, possibly uncovering a generalised algorithm to update any representation.

## Introduction

The brain is constantly bombarded with all types of information that need to be related to each other in order to build a coherent picture of one’s surroundings (or of the situation one is experiencing at a given time). A broad range of studies have led to assert that one of the main roles of the hippocampal region is to do that: integrate multimodal information – coming from a variety of sensory and associative cortices, as well as deeper structures – to build a dynamic representation of an environment or an event ^1–4^. The accurate updating of these representations – and their comparison with previously stored representations as well as projections of likely near futures – could allow one to evaluate the outcome of a range of decisions in order to react adequately. In the context of spatial cognition, information integration is implemented in interconnected subareas of the hippocampal region through neurons coding for specific instantaneous navigational features such as position (place cells) ^5^, direction (head direction cells) ^6^, local metrics (grid cells) ^7^ and boundaries (border cells) ^8, 9^. Successful navigation is also thought to depend on the accurate updating of spatial representations, which themselves would hinge on self-motion signals and their integration with both positional and directional information ^10–13^. Despite their crucial role, where and how self-motion signals are integrated remains largely elusive.

Linear speed modulation has so far mainly been reported in the CA1 region of the hippocampus and the medial entorhinal cortex (MEC) in conjunction with positional information or as an (apparently) self-standing code (speed cells) ^14, 15^. In addition, speed has been reported to influence oscillatory activity recorded in the hippocampal field potential where the theta power seems correlated to locomotory activity ^16–18^. In contrast, angular velocity coding has not been yet established in principal neurons of the hippocampal region. Most reports come from recordings of subcortical structures (e.g., lateral mammillary nuclei, dorsal tegmental nucleus), linked to the processing of vestibular information ^19–22^.

Most crucially, it has remained unclear whether angular velocity coding reflects an intelligent design. In fact, some theoretical models within the class of continuous attractor networks (CANs) require that neurons coding for instantaneous navigational correlates – such as position and direction – receive inputs from cells conjunctively coding for position, direction and self-motion ^11, 13, 23–27^. These conjunctive cells – sometimes refers to as the ‘hidden layer’ – are hypothesised to mediate the shift of activity from position (or direction) at time *t* to the next position (or direction) at time *t+1* (Figs. 1c and 2b). Exciting evidence compatible with such CAN models was recently provided by investigations of the *Drosophila melanogaster* central complex, where head direction cells whose activity was modulated by angular velocity were shown to be organised in a ring according to their preferred direction ^28, 29^. These reports and the underlying mechanisms are likely restricted, in insects, to directional coding.

**Fig 1.**
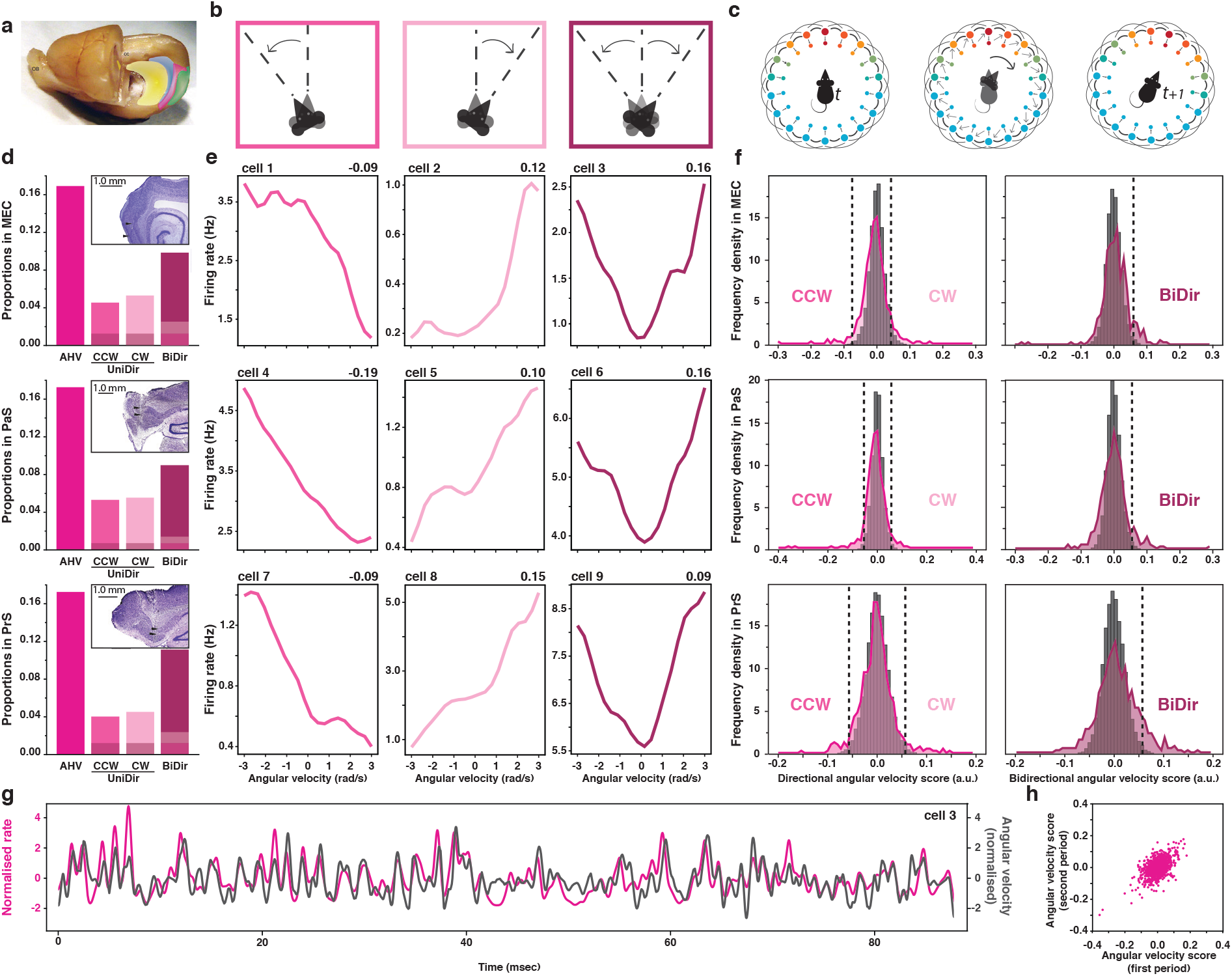
Angular velocity cell in the parahippocampal cortex. (**a**) Whole rat brain, with partially removed left hemisphere to enable a midsagittal view of the right hemisphere and outlines of hippocampal formation (yellow), presubiculum (blue), parasubiculum (pink) and MEC (green). Adapted from ^30^ (**b**) Schematic representation of three type of angular head velocity (AHV) movement, from left to right: counterclockwise (CCW, dark pink), clockwise (CW, light pink) and bidirectional (BiDir, purple). (**c**) Schematics ring attractor depicting theoretical updating of head direction code from position at time_*t*_ (left) to position at a later time_*t+1*_ (right) following angular movement (middle). The outer layer of head direction (HD) cells is connected to a “hidden” inner layer of conjunctive HD-by-AHV cells. The colour represents neural activation from maximum (red) to minimum (blue). (**d**) Proportions of AHV cells within MEC (top), PaS (middle) and PrS (bottom). From left to right: all AHV cells (fuchsia, MEC:17%, n=67; PaS:17%, n=75; PrS:17%, n=104), CCW-AHV cells (dark pink, MEC:5%; PaS: 5%; PrS:4%), CW-AHV cells (light pink, MEC:5%; PaS: 6%; PrS: 5%), and BiDir-AHV cells (purple, MEC:10%; PaS:9%; PrS:11%). The shaded areas represent the intersection between AHV-CCW & AHV-BiDir (darker shade) or between AHV-CW & AHV-BiDir (lighter shade). Upper right corner boxes: representative Nissl-stained sagittal section showing example recorded track for each area. (**e**) Example AHV cells in MEC (top), PaS (middle) and PrS (bottom) showing firing rate as a distribution of angular velocity (in rad/s), score in upper right corner. From left to right: CW-AHV (dark pink), CCW-AHV (light pink) and BiDir-AHV (purple). (**f**) Distribution of unidirectional (left) and bidirectional (right) AHV scores across MEC (top), PaS (middle) and PrS (bottom) cell population comparing observed (coloured curve) and shuffled data (grey bars). Dashed lines represent 99 percentile thresholds for CCW- and CW-AHV (left) and Bidir-AHV (right). (**g**) Snapshot comparison between z-scored firing rate of an example AHV cell (pink curve) and instantaneous angular head velocity (black curve). (**h**) Correlation between the AHV scores calculated in the first and second half of each recording sessions (rho stability = 0.52).

**Fig 2.**
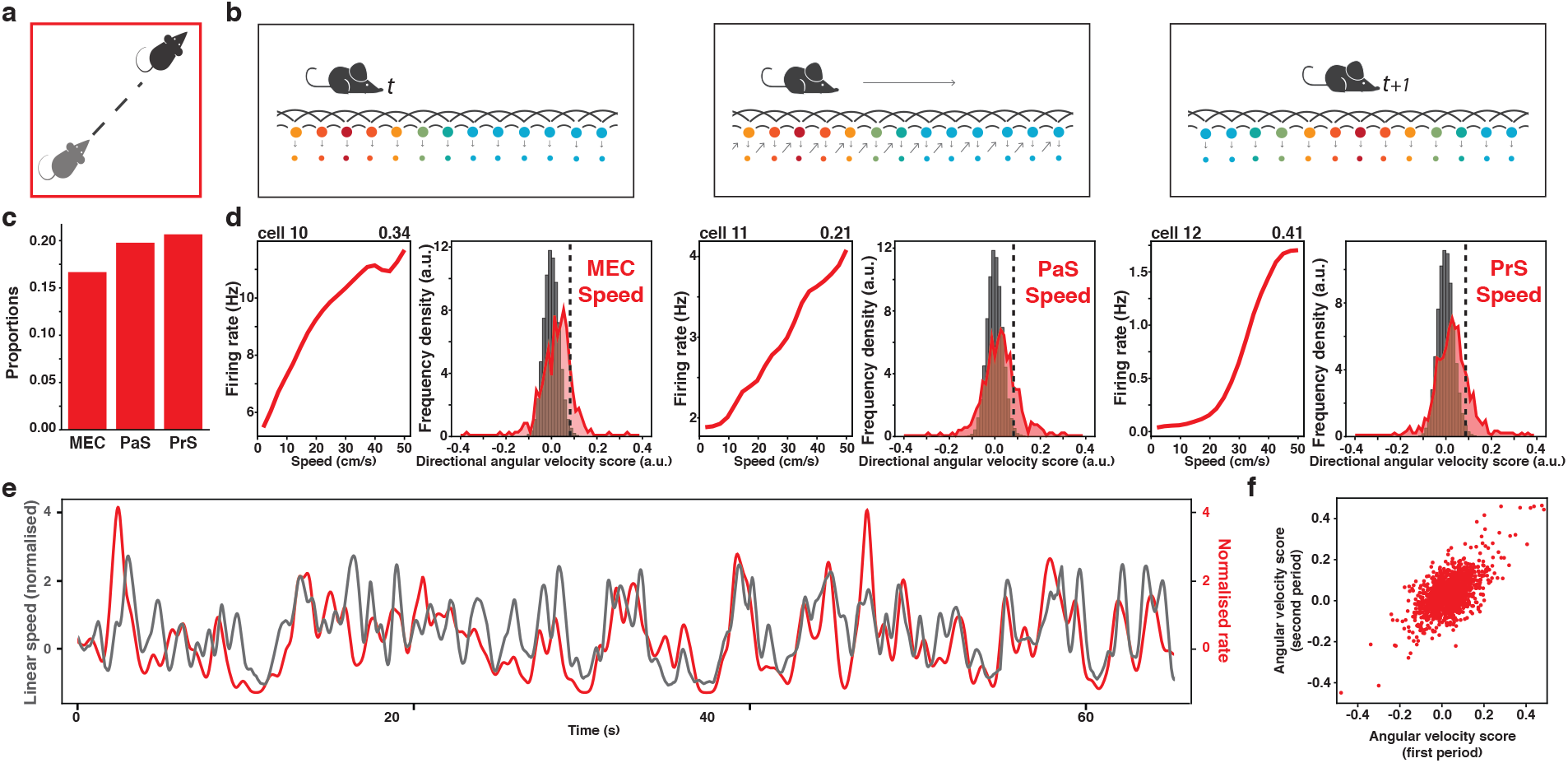
Speed cells are distributed across the parahippocampal cortex. (**a**) Schematic representation of linear velocity movement (**b**) Schematics partial ring attractor depicting theoretical updating of positional code from position at time_*t*_ (left) to position at a later time_*t+1*_ (right) following linear movement (middle). The outer layer of conjunctive grid-by-HD cells is connected to a “hidden” inner layer of conjunctive grid-by-HD-by-speed cells. The colour represents neural activation from maximum (red) to minimum (blue). (**c**) Proportions of speed cells in MEC (left, 16.7%, n=66), PaS (middle, 19.8%, n=86) and PrS (right, 20.1%, n=125). (**d**) Example speed cells (left) and general distribution of speed scores (right) in MEC (left panel), PaS (middle panel) and PrS (right panel). The example tuning curve on the left plot shows the mean firing rate (red) as a function of speed, between 2 cm/s and 50 cm/s. Scores are in the upper right corner. The histograms on the right compare the observed (coloured curve) and shuffle data (grey bars), dashed lines represent the 99-percentile threshold. (**e**) Snapshot comparison between z-scored firing rate of an example AHV cell (red curve) and instantaneous linear speed (black curve). (**f**) Correlation between the speed score calculated in the first and second half of the recording session (rho stability = 0.61).

To understand the circuit mechanism by which spatial representations can be updated in mammal, we carefully mapped the activity of individual neurons recorded in three main areas of the rat parahippocampal region: the MEC, the parasubiculum and the presubiculum. We specifically investigated whether these neurons could respond to both linear and angular self-motions signals. Our study revealed the existence of parahippocampal neurons coding conjunctively for direction, position and self-motion, possible evidence for the elusive “hidden layer”, pillar of many CAN models. However, the thorough examination of self-motion neurons exposed several inconsistencies with such models, highlighting a need for their revision. In particular, it appears that direction, position and speed selectivity are randomly admixed with each other, perhaps pointing to a capacity for self-organization of these circuits. Given that linear and angular speed are the derivative in time of position and direction respectively, this may reflect a general self-organizing derivative algorithm for the updating of any type of information.

## Results

To understand whether and how both linear and angular speed modulation are integrated in the hippocampal circuits, we analysed the activity of 1436 principal neurons recorded in all layers of three interconnected subareas of the parahippocampal region of rats freely exploring open environments of various sizes (MEC: 396 cells; presubiculum: 605 cells; parasubiculum: 435 cells, Fig. 1a) ^30^.

### Angular velocity coding in the parahippocampal region

First, we sought to determine whether parahippocampal neurons responded to the angular head velocity (AHV) signal, which is the derivative of head direction in time. To that end, we computed, for each cell, its angular velocity score as the Pearson product moment correlation between the instantaneous value of angular velocity and the firing rate of the cell across the recording session (see Methods and extended data Fig. 4a). We defined cells as AHV modulated when their score was greater than the 99th percentile of the shuffled distribution. This method led us to classify a total of 246 cells as angular head velocity cells, amounting to about one sixth of all parahippocampal cells (MEC: 16.9%; Prs: 17%; PaS: 17.2%; Figs. 1b–e and extended data Figs. 1a–d). AHV modulation was uniformly distributed across all layers of each region, except in MEC LII which showed no such modulation (Kolmogorov Smirnov test, p< 0.001, extended data Figs. 2a–c).

As per our definition, AHV cells are neurons whose firing rate is positively modulated by angular velocity, meaning that these cells are more active when the animal is turning its head. About half of the AHV cells had their activity modulated solely when the animal had its head turning only in one direction, either clockwise (CW) or counterclockwise (CCW). The other half of the AHV cells were bidirectional (BiDir) and modulated by angular motions in both directions (MEC: 26.8% CW, 31.0% CCW, 58.2% BiDir; Prs: 25.9% CW, 23.1% CCW, 64.4% BiDir; PaS: 32.0% CW, 30.0% CCW, 52.0% BiDir; Figs. 1b–f and extended data Figs. 1a–d, note that the percentages do not sum to 100%, since unidirectional and bidirectional scores are not mutually exclusive, see Methods). All layers of each region presented similar proportions of CW, CCW and bidirectional AHV cells (Kolmogorov Smirnov test, p< 0.001, extended data Figs. 2a–c) with naturally the exception of MEC layer II. AHV modulation appeared stable in time across all regions and we observed no change in modulation intensity (AHV score) while comparing successive half-sessions (Pearson correlation ρ=0.52, p< 0.001, Fig. 1h).

To reduce the dependence of our results on a unique scoring method, we further analysed all recorded cells with two additional methods. Given that the Pearson correlation would present higher scores for linear modulation, we selected methods that would allow us, for example, to select cells responding to a specific speed band. The first additional score was inspired by a method commonly used to characterized spatial modulation ^31^. The “Skaggs score” of each cell was obtained by computing the information per spike that each neuron conveyed about the running speed or AHV (see Methods). The second additional method relied on a generalized linear model (GLM) approach ^14^ to calculate a neuron firing profile as a function of velocity values. For this approach, velocity values were treated as categorical variables; therefore, no linear dependence was implied (see Methods and extended data Fig. 4b). While these additional methods revealed a substantial fraction of AHV modulated cells not captured by linear scoring method (and vice-versa), the cell populations yielded by the three methods were significantly overlapping (Binomial test, p < 0.001, extended data Fig. 4c). Out of 182 AHV cells solely picked up by the GLM method, about half of them (54%) were anticorrelated with the absolute value of angular velocity – meaning that their activity was maximal when the animal was not turning its head. Only a small number of them (11%) were responsive to a particular value of angular head velocity (measured with a gaussian fit), while the rest did not show a modulation profile with a simple shape. Given the dominance of linear coding and to facilitate the comparison with linear speed analyses, we decided to use the more conservative Pearson scoring methods for all further AHV analyses.

### Parahippocampal neurons upstream of the entorhinal cortex code for linear speed

Once established that angular velocity coding was widespread across several parahippocampal areas, we tested whether linear speed coding could also extend beyond the medial entorhinal cortex (MEC). To that end, we determined the speed score of each cell as a Pearson product moment correlation between the instantaneous value of rectilinear speed and the firing rate of the cells across the recording session (see Methods and extended data Fig. 4a). With this method, we classified a total of 277 speed cells. Our results confirmed the existence of speed cells uniformly in all layers of the MEC in similar proportions to what was previously reported ^15^ (MEC all: 16.7%, LII: 23.9%, LIII: 18.7%, LV: 12.7% and LVI: 13.7%; Figs. 2a–e and extended data Figs. 1a–b and 2a). In addition, we observed that rectilinear speed signals could be found upstream of the MEC, in about one fifth of both PrS and PaS cells (Prs: 20.6%; PaS: 19,8%, Figs. 2a–e and extended data Figs. 1c–d). Speed cells were uniformly distributed across all layers in each area (Kolmogorov-Smirnov test, pvalue<0.001; extended data Figs. 2a–c). As for AHV cells, speed cells were stable across time (Fig. 2f).

We further analysed speed modulation with two “non-linear biased” methods (i.e., “Skaggs information per spike” and GLM) similar to those we used for AHV cells. The population of speed cells detected with these methods significantly overlapped with those obtained with our linear scoring method (binomial test, pvalue < 0.001; extended data for GLM in Fig. 4c). Similar to AHV scoring, the Pearson method yielded the lowest number of speed-modulated neurons (Skaggs and GLM approaches respectively resulted in 334 and 551 selected neurons). Out of 296 speed-modulated cells solely picked up by the GLM method, about one fifth were anticorrelated with the absolute value of speed and about one sixth showed a very low linear correlation. Only a very small number of them (8%) were responsive to a particular speed band (measured with a gaussian fit), while the rest did not show a modulation profile with a simple shape. In order to allow for comparison with the latest reports of speed cells in the MEC, we decided to use the more conservative Pearson scoring methods for all further analyses.

### Conjunctive coding of primary (place, direction) and derivative (velocity) signals

Based on the fact that angular head velocity (AHV) and speed are the derivative in time of head direction and position respectively, we defined positional and directional signals as primary and self-motion signals as derivative.

Given the key role of a “hidden layer” of cells presenting conjunctive coding for continuous attractor network (CAN) theories ^11, 32^, we next sought to determine to which degree self-motion signals are co-existing in conjunction with other types of coding at the unit level. To that end, we computed the grid and head direction (HD) scores of each recorded unit. We labelled as ‘significantly modulated’, cells whose score exceeded the 99th percentile of the score distribution calculated on shuffled data (see Methods). According to these parameters, the majority (80.4%) of AHV cells coded for at least one other feature (HD: 55.2%; grid: 13.4%; grid-by-HD: 5.6%; rectilinear speed: 33.7%; Figs. 3a–b, 3e and 4a–c). These percentages were similar to the percentages observed in the general population (HD: 53.3%; grid: 17.8%; grid-by-HD: 7.7%, rectilinear speed: 19,2%; Fig. 3a). A similar distribution of conjunctive coding was observed among speed cells (HD: 35.4%; grid: 18.7%; grid-by-HD: 3.2%; AHV: 29.9%; Figs. 3c–d, 3f and 4a–c). We observed all possible types of conjunction of code including AHV-by-HD (the hidden layer of directional CAN models) and grid-by-HD-by-speed (the hidden layer of positional CAN models). This result contrasted significantly with previously published studies of a predominantly self-standing code in speed cells recorded in the MEC superficial layers ^15^.

**Fig 3.**
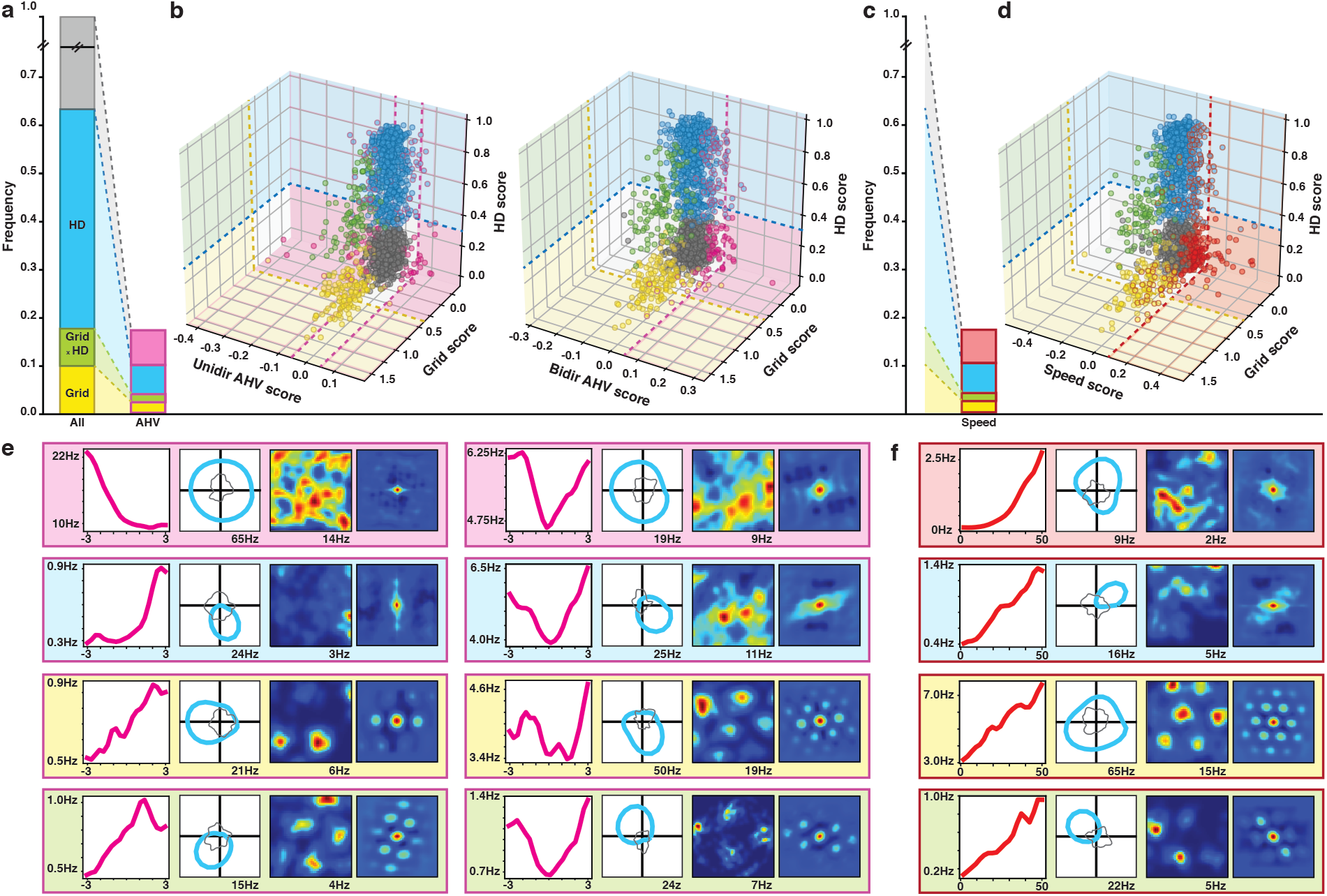
The majority of speed and angular velocity cells presents conjunctive coding. (**a**) Intersection between angular head velocity (AHV), grid and head direction (HD) cells. The bar on the left shows the proportions of cells qualifying as grid (yellow, 17.8%), HD (blue, 53.3%) and grid x HD (green, 7.7%) cells in the total population. In grey are cells neither coding for HD nor grid. The bar on the right (pink contour) shows the same distribution among cells qualifying as AHV cells (grid: 13.4%; HD: 55.3%; grid x HD: 5.7%). (**b**) Scatterplots showing the intersection between grid, HD and AHV scores. Plot on the left: unidirectional – UniDir – AHV: i.e., CCW-AHV and CW-AHV. Plot on the right: bidirectional – BiDir– AHV. The colour code is the same as in (a). A pink contour denotes a modulation by AHV. Dotted lines represent region-averaged classification thresholds, computed to guide the visualization. (**c**) Same as in (a) for speed cells. The red contour bar represents percentages of principal cells within the speed cell population (grid: 18.8%; HD: 35.4%; grid x HD: 3.3%). (**d**) Same as in (b) for speed cells. A red contour denotes a modulation by speed. (**e**) Examples of the four different kinds of AHV modulated cells, colour coded as in (a). From left to right: AHV tuning curve, HD polar plot, spatial firing rate map, spatial autocorrelogram. (**f**) Examples of the four different kinds of speed modulated cells, colour coded as in (c). Same plot as in (e).

**Fig 4.**
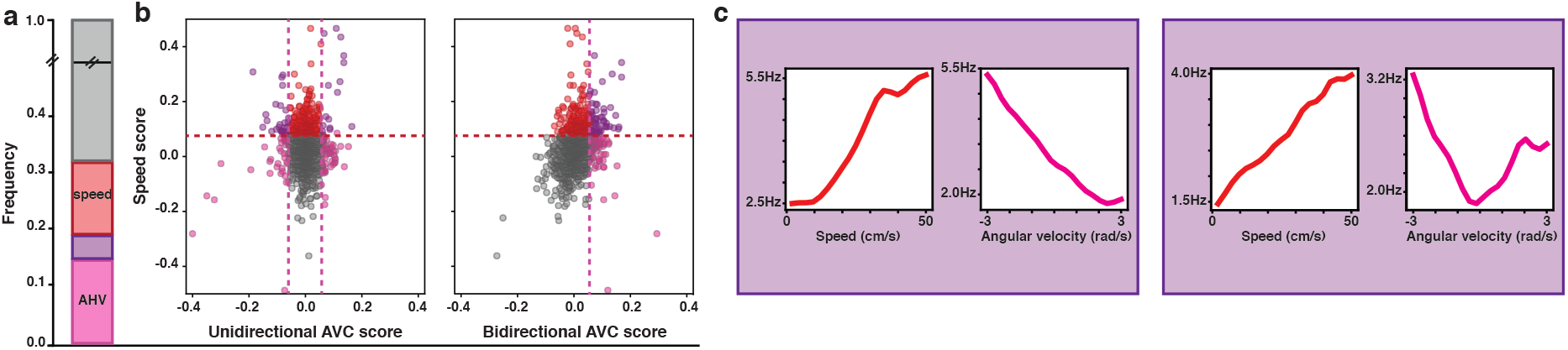
Intersection between angular head velocity and speed cells. (**a**) The bar shows the percentages of cells in the whole population. Pink: cells coding only for AHV (11.3%); Red: cells coding only for speed (13.5%); Purple: cells coding for speed and AHV (5.8%); Grey: cells neither coding for AHV nor speed. (**b**) The scatterplots show the intersection between AHV and speed scores in the population. Plot on the left: unidirectional UniDir – AHV: i.e., CCW-AHV and CW-AHV. Plot on the right: bidirectional – BiDir– AHV. Colour code as in (a). Dotted lines represent region-averaged classification thresholds. (**c**) Examples of two kind of conjunctive AHV X speed cells. Plots show the firing rate as a function of the angular velocity (pink) or speed (red). Left panel: speed x UniDir-AHV cell. Right panel: speed x BiDir-AHV cell.

To test whether these differences could be explained by regional variations and an over-representation of MEC LII cells in previous reports, we compared the percentage of conjunctive cells across all layers of MEC, presubiculum and parasubiculum. The percentage of conjunctive cells was similar in all layers (extended data Fig 5a-c), with the exception of MEC LII, where the percentage of cells coding conjunctively for a primary and a derivative signal (3%) was significantly lower than the one in the general population (18%, t-test, pvalue <0.05). The scores (grid, HD, AHV and linear speed score) were mostly independent from each other and we did not observe any significant correlation between them, with the exception of a relatively small correlation between speed and bidirectional AHV scores (Pearson r= 0.29, pvalue < 0.001) and a small anticorrelation between speed and HD scores (Pearson r=-0.14, pvalue <0.001). The observed distribution of mixed selectivity is compatible with a simple hypothesis of independent assignment of each of the coding properties in the general population: cells coding for different behavioural features neither segregate, nor cluster together. This was true for all conjunction combinations and all layers, except for PrS deep layers and MEC LIII, in which an under representation of speed x HD cells was found (binomial test, pvalue <0.05 for MEC LIII, pvalue < 0.01 for PrS deep). That self-motion information is integrated at the unit level in all cell types and all tested layers (with the notable exception of MEC LII) is incompatible with current CAN model and calls for their revision.

### Derivative signals seemed encoded differently than primary signals

In order to grasp whether derivative signals (i.e., speed and AHV modulation) were encoded in a similar fashion as primary signals (i.e., position and direction modulation), we compared the firing properties of each class of neurons. We observed that cells coding for derivative signals exhibited higher average firing rates than cells coding for primary signals (t-test: pvalue < 0.001; extended data Fig. 6a). They also showed a shorter average inter-spike interval (t-test: pvalue <0.001; extended data Fig. 6c) and a larger peak firing (defined as the fifth quintile of the rate distribution, t-test: pvalue <0.001; extended data Fig. 6b). It is important to notice that, contrary to what would be expected from CAN models, unidirectional AHV cells were not silent when the rats were not turning their head. Instead, they decrease their rate in response to movement in one direction and increase it in the other. The differences in firing properties between primary and derivative neurons could be explained by the fact that the monotonic firing profiles used to encode motion signals is less sparse than the receptive-field coding of grid and HD cells which are largely silent outside their firing field. To test this hypothesis, we calculated the percentage of the correlate values at which each cell fired more than its average firing rate (see Methods and extended data Fig 6d). Derivative cells showed a significantly larger percentage (mean: 61.3%, std: 15%) than primary cells (mean: 29.8%, std: 9.4%; t-test: pvalue < 0.001).

To further characterize whether derivative signals are monotonically encoded in the parahippocampal region, we fitted the rate-response tuning curve of both AHV- and speed modulated neurons with either a linear or a sigmoid function (see Methods). The majority of AHV cells (68 %) were better described by a linear fit, compared to a sigmoidal fit (extended data Figs. 3a–c). As the steepness parameter of the sigmoidal fits was generally low (mean: 0.47 (rad/s)^-1^, std: 0.16 (rad/s)^-1^), we concluded that most AHV cells followed a quasi-linear rate function. In contrast with the AHV population, the sigmoidal fit with low steepness was slightly more predominant among speed cells (56%). This result seemed to be due to a saturation observed at high speeds and was probably related to low sampling in that speed band (extended data Figs. 3a–c). Yet, also in this case, the fitted sigmoids generally exhibited low steepness (mean: 0.05 (cm/s)^-1^, std: 0.025 (cm/s)^-1^). We thus concluded that most speed cells also followed a quasi-linear rate function.

### Velocity coding is independent of theta modulation

Because the theta rhythm of the local field potential has historically been strongly associated to running speed, we investigated whether both AHV and speed cells had their activity modulated by theta. Following previous work ^30^, we defined that a cell is theta modulated when its mean spectral power around the peak in the 5-11 Hz range was at least fivefold greater than the average spectral power in the 0-125 Hz range (see Methods). We observed that only around 40% of the AHV and the speed cells passed these criteria for theta modulation (Fig. 5a), while many of the remaining cells did not show any modulation by theta (Fig. 5b–c). The theta modulation of AHV/speed cells was observed in comparable proportions to those observed in the general population (Fig. 5e). There was no significant correlation between AHV score and theta score, or between speed score and theta score (Pearson correlation: pvalue > 0.05). Conjunctive coding for grid or head direction did not influence the proportions of velocity cells that were theta modulated (Fig. 5f, t-test: pvalue < 0.05). Theta modulation was uniformly distributed across all layers of each area except for MEC LII which showed more theta modulation and MEC LVI which showed less (extended data Fig. 7a, t-test: pvalue <0.01). The proportion of velocity cells theta modulated in each layer did not differ from what was expected based on the theta modulation observed in the general population, except for MEC LVI speed cells, which showed less theta modulation than expected (extended data Fig. 7b–c, t-test: pvalue <0.05). Together, these results suggest that the code for self-motion in the parahippocampal region seems largely independent of theta modulation at the single cell level.

**Fig 5.**
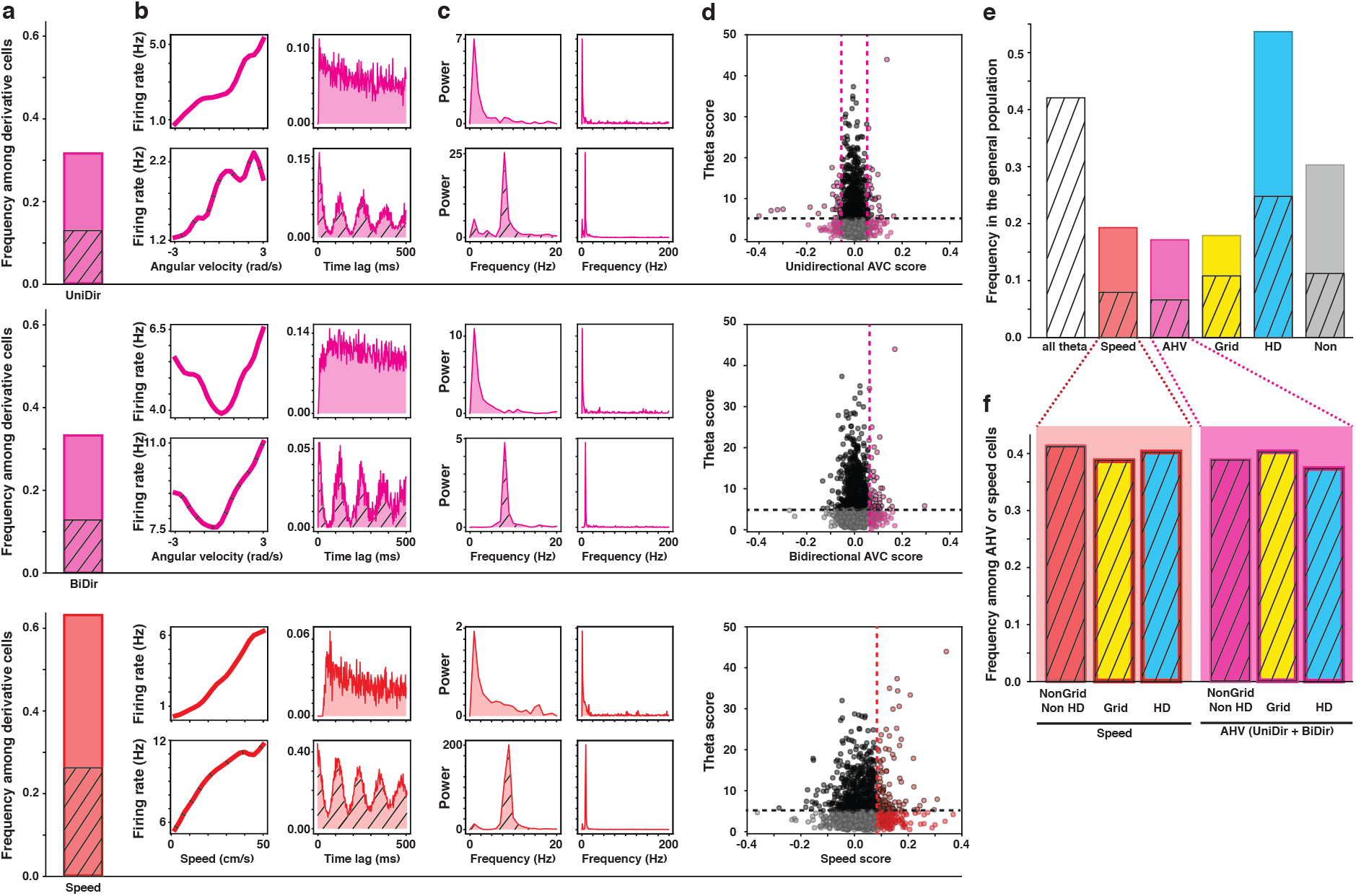
Speed and angular velocity coding are independent from theta modulation. (**a**) Frequency of unidirectional AHV cells (first row, 31.1%, n=137), bidirectional AHV cells (second row, 32.9%, n=145) and speed cells (third row, 62.9%, n=277) within the population of derivative cells. Dashed bars represent the fraction of theta modulated cells. UniDir-AHV: 12.5%, n=55; BiDir-AHV: 12.2%, n=54; speed: 25.9%, n=114. (**b**) Examples of theta modulated (bottom row of each section) and non-modulated (top row each section) derivative cells. The first column shows the firing rate as a function of the angular velocity or speed, the second column shows the time autocorrelogram of the firing rate of the cell. (**c**) Power spectrum of cells in (b). First column: power in the range 0-20 Hz; second column: full spectrum (0-200 Hz) (**d**) Scatterplots of the distribution of theta scores and unidirectional AHV scores (first row), bidirectional AHV scores (second row) and speed scores (third row) in the population. Full black dots represent cell that are theta-modulated but not modulated by AHV or speed; Full coloured dots represent cells modulated by AHV (pink) or speed (red), but not by theta; coloured dots with black contour are cells modulated by both theta and AHV (pink) or speed (red); grey dots represent unclassified cells. (**e**) Distribution of speed cells (red, 19.3%), AHV cells (pink, 17.1%), grid cells (yellow, 17.8%), HD cells (blue, 53.3%) and unclassified cells (grey, 30.0%, n=432) in the whole population. Dashed bars represent the frequency of theta modulated cells in the whole population (white shaded, 42.1%), and in conjunction with speed cells (8.0%), AHV cells (6.6%), grid cells (10.1%), HD cells (24.6%) and unclassified cells (11.1%). (**f**) Frequency of theta modulated cells among speed cells (left, red background) and AHV cells (right, pink background). Cells are divided by type: “pure speed” (red, 38.2%), “pure AHV” (pink, 28.6%), conjunctive grid x speed (yellow left, 51.9%), conjunctive grid x AHV (yellow right, 60.6%), conjunctive HD x speed (blue left, 41.8%) and conjunctive HD x AHV (blue right, 43.4%).

## Discussion

Here we reveal the existence of a network of parahippocampal principal neurons whose activity is modulated by both angular and linear self-motion signals. Extensive mapping of the rat parahippocampal region showed that this network was spread homogenously across all layers of several interconnected areas upstream of the hippocampal formation: the medial entorhinal cortex (MEC), the presubiculum (PrS) and the parasubiculum (PaS). We observed that some neurons modulated by self-motion seemed to only respond to either angular head velocity or linear speed (i.e., an apparent self-standing code). Yet, the majority of those self-motion neurons were also concomitantly responding to spatial or directional information (i.e., a conjunctive code). Such integration at the unit level may be a crucial mechanism underlying the generation and the updating of the representation of position (place and grid cells) and direction (head direction cells) in the hippocampal/parahippocampal circuits. Moreover, given that linear and angular speed are the derivative in time of position and direction respectively, our results may uncover a general algorithm for the updating of any type of information.

In support of this idea of a general parahippocampal mechanism that could compute the derivative in time of other correlates, we demonstrated that both angular and linear self-motion (derivative) signals were encoded in a different manner with respect to positional and directional (primary) signals. It is well documented that head direction, grid, and place cells tend to be active only when an individual is either in a given position or with its head in a given direction. In contrast, we showed that only a negligible proportion of self-motion neurons responded preferentially for a given speed. The vast majority presented a much higher baseline activity and had their firing rate linearly (or quasi-linearly) ramped up in response to increasing speed. This could be the hallmark of a general strategy in the parahippocampal region for the neural coding of scalar quantities whose magnitude has a well-defined meaning – speed and angular velocity in this case – as opposed to neural activity manifolds used to encode position and direction, where coordinates are always relative to a reference frame.

Previously, angular head velocity (AHV) cells had been mainly characterized upstream of the hippocampal circuits, in subcortical structures linked to the processing of vestibular information (e.g., lateral mammillary nuclei, dorsal tegmental nucleus, thalamic nuclei and striatum) ^19–22, 33, 34^. In addition, some example of AHV modulation was reported in the retrosplenial cortex ^35, 36^ and has been linked to the accurate processing of visual inputs in the primary visual cortex ^37^. Besides, a few examples of hippocampal neurons modulated by whole body motion have been reported in the primate ^38^. Furthermore, angular head velocity has been shown to influence the preferred orientation or the selectivity for pitch and azimuth of some presubicular head direction cells ^36, 37, 39^, as well as to modulate the firing rate of a small fraction of fast spiking presubicular interneurons ^40^. We observed both bilateral and unilateral (i.e., CW and CCW) AHV modulation in the presubiculum, the parasubiculum and the MEC. We hypothesise that such signals were not reported in previous studies either because of a restricted scope in analyses or because recordings were often clustered in the most dorsolateral part of the presubiculum, at the border of the retrosplenial cortex. To avoid such bias, we recorded from a much larger population spread out across the medio-lateral and antero-posterior axis. Nevertheless, it is worth noting that our recordings did not show any topographic organisation of self-motion modulation. The lack of topography, in areas near the top of the cortical hierarchy is consistent with AHV being one of potentially several high-level signals, which could be extracted by derivation with respect to time ^41^.

Since the discovery of grid cells, many have attempted to understand how such a strikingly regular signal could be generated by individual neurons ^11, 23, 42–45^. A speed code is central to much of this theoretical work, and a break-through was the characterization of speed cells and speed modulation in the MEC ^14, 15, 46–49^. Interestingly, recent experimental work has demonstrated that grid cell activity is dependent on the integrity of the speed signal^50^, which itself seems driven by the brainstem locomotor circuit ^51^. Likewise, the stability in head direction coding seems dependent on angular head velocity (AHV) signal ^52^ and vestibular inputs ^20, 53^. Therefore, self-motion signals could be similarly involved in the generation and the maintenance of both position and direction signals. It is interesting to note that MEC LII contains neither HD nor AHV signals, a major exception to the apparent random assignation of correlates, arguing for the importance of a local network in some cases.

Our findings offer robust experimental evidence particularly relevant for the reappraisal of grid and head direction cell generation theories based on continuous attractor neural networks (CAN). CAN are popular theoretical models of how reciprocally interconnected neurons may extract, refine and sustain stable representations of continuous behavioural variables. This function is particularly crucial when afferent signals are weak, noisy and/or intermittent. How can such representations be updated? A common view is that neurons coding for “primary” navigational variables – such as position and direction – are connected by a so-called “hidden layer” of cells conjunctively coding for position, direction and self-motion. This conjunctive layer is postulated to drive the activity of the output layer to remain congruent, at any time, with the external world. Although several variants of these mechanisms have been proposed, in general they posit a categorical division of labour between the output units representing the instantaneous variables and the hidden units updating them ^23, 54^. Recent studies in dr*osophila melanogaster* have indeed shown that a ring attractor network with local excitation and global inhibition underlies the representation of head direction ^28, 29^. It is possible that similar mechanisms operate in mammals. Yet, it is difficult to imagine how a similar neural structure, neatly laid out and possibly hard-wired in the fly brain, could be engineered in the absence of topography in the mammalian brain, and be retained in areas with such prominent plasticity.

Here we report the existence of cells conjunctively coding for position, direction and self-motion in both the pre- and the parasubiculum as well as in the medial entorhinal cortex, providing the first recording in the hippocampal region of the conjunctive coding postulated to be found in the “hidden layer” of CAN models. Yet, our findings should not be perceived as a blanket endorsement of such models. Indeed, our results challenge theories assuming very specific architectures, such as those requiring that *all* speed and AHV cells must conjunctively code for position and head direction. While we observed more conjunctive motion-by-primary signal coding than what was reported in previous studies of speed cells in MEC layer II ^15, 47, 49^, we still observed a significant number of self-standing AHV and speed cells. Furthermore, the distinct types of selectivity not only appeared randomly admixed, but also did not present evidence of clear-cut categorical distinctions between conjunctive and non-conjunctive cells. We observed that scores (i.e., grid, HD, AHV and linear speed score) showed no correlation, and the observed percentages of conjunctive cells were compatible with a scenario of independent assignment of coding properties. Such results are thus more consistent with self-organizing models than with precisely engineered ones ^45, 55^. Ultimately, they challenge the very concept of a well-defined representation of the instantaneous variables. In terms of attractor structures, they support extending CAN models to incorporate dynamical variables in what their output “represents”, for example including units firing at a baseline rate for zero AHV, and quiescent for high either CW or CCW AHV. However, theoretical work on such dynamical attractors or “structured flow manifolds” is only just starting ^56, 57^. It will have to take into account the remarkable lack of segregation of the parahippocampal regions in the representation of different correlates. One exception to this observation was the absence of AHV and HD cells in MEC LII, suggesting that the AHV signal is needed locally, among the same cells coding for the primary signal it serves to update. In the framework of attractor network models, this leads us to hypothesize that the neural circuitry representing and updating dynamical variables is one and the same, not only at the level of the brain area, but also locally within a layer and even within individual cells. Interventional studies will be necessary to test how such randomly assorted conjunctive coding of primary and self-motion signals can be necessary for accurate spatial cognition.

Historically, running speed has been reported to show a strong correlation with the amplitude of theta oscillations recorded in the local field potential of freely behaving rodents ^16–18^. Likewise, many place and grid cells exhibit a strong modulation of their firing rate following those theta oscillations, either in a phase-locked or in a phase-precessing manner ^58–60^. In line with these observations, many models point to theta oscillations as inherent to the generation of the grid signal. Among them, the oscillatory interference-based models of grid cells assume a velocity input to the grid network composed of translational speed and movement direction ^42, 44^. Here we show that only 40% of our speed cells and AHV cells show a strong modulation by theta, a finding that should be taken into account in refined versions of such models. This apparent decoupling between speed/velocity signals and theta may be surprising for some. However, the role of theta in spatial coding had already been recently challenged by two lines of evidence. One is that modulation of the septal oscillatory activity had no apparent consequence on grid signal maintenance ^61^, therefore suggesting that non-theta septum correlates – such as attention – were involved in the grid cell signal disruption observed after septal inactivation ^62, 63^. Second is that no stable theta oscillation had been recorded in the hippocampal local field potentials of bats, who do exhibit both place and grid coding^64^. Yet, recent results show that bats seem to exhibit theta-band modulation of grid firing and matching phase precession when considering a non-stable theta frequency ^65^. Further targeted interventional studies would be essential to determine the ambiguous role of theta in the generation and maintenance of the grid signal. Discriminating between the theta modulated and the non-theta modulated self-motion neurons, revealed by our work, would be a chief target for these studies.

In conclusion, we provide clear evidence of a widespread parahippocampal network involved in linear and angular speed coding that could have a crucial role in the updating of the cognitive map, or perhaps be part of the map itself. The existence in the hippocampal region of neurons conjunctively coding for self-motion, position and direction would *prima facie* appear to fill a gap in the framework of continuous attractor network models. Yet, the analysis of these neurons reveals features divergent from those expected of units serving solely to update the instantaneous representation of static variables. Our work urges for the revision of such models so that they can (i) express dynamical continuous attractors and (ii) account for the apparent random nature of the spatial code, as well as its peculiar lack of a clear organization. We hypothesize that derivative algorithms may have a generalized role in the updating of continuously varying information, not just of a spatial nature. Further studies, with either targeted inactivation of neurons or testing of non-spatial correlates will be necessary to establish whether one of the main roles of the parahippocampal region is to ensure the accurate updating of the hippocampal representation.

## Acknowledgement

We thank Edvard and May-Britt Moser for consent to use of these data ^30^. We thank René Huster and Arnfinn Aamodt, students in the laboratory of Johan Frederik Storm, for preliminary analyses of head direction signal. We thank Silvia Girardi for her help with the schematics in Figs. 1 and 2 illustrating ring attractors. We thank Dori Derdikman, Gily Ginosar, Eleonore Duvelle and Anna Chambers for providing comments on an earlier version of the manuscript. Charlotte Boccara is supported by a MSCA fellowship and an NFR grant.

## Contributions

C.N.B. conceptualized the experiments. C.N.B. and D.S. conceptualized the analyses. D.S. perform the analyses with support by C.N.B. and A.T. C.N.B. and D.S. wrote the paper with feedback from A.T.

## Extended data

**SF1.**
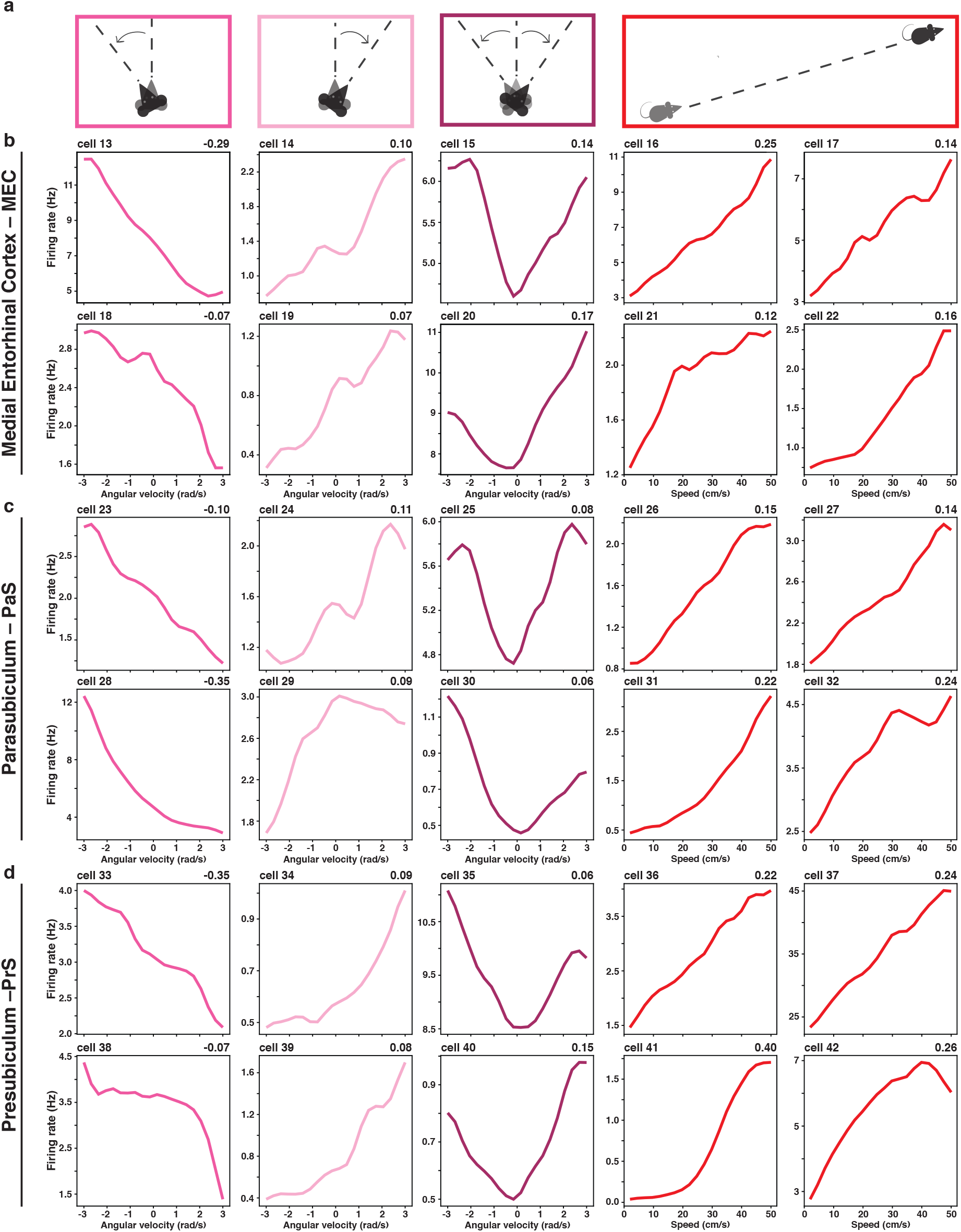
Extended examples of speed and Angular head velocity cells. (**a**) Schematic representation of three type of angular head velocity (AHV) movement and linear speed, from left to right: counterclockwise (CCW, dark pink), clockwise (CW, light pink), bidirectional (BiDir, purple) and linear speed (red). ^30^ additional examples of self-motion cells: 6 AHV and 4 speed cells in each region; medial entorhinal cortex (**b**), parasubiculum (**c**) and presubiculum (**d**). The firing rate is represented as a function of angular velocity (in rad/s) or speed (in cm/s). AHV or speed scores are reported in the upper right corner. Cell ID are reported in the upper left corner. From left to right: CCW-AHV (dark pink), CW-AHV (light pink), BiDir-AHV (purple) and speed (red, last two columns). Note that the high values of the rate of cell ^37^ suggest that it may be an interneuron.

**SF2.**
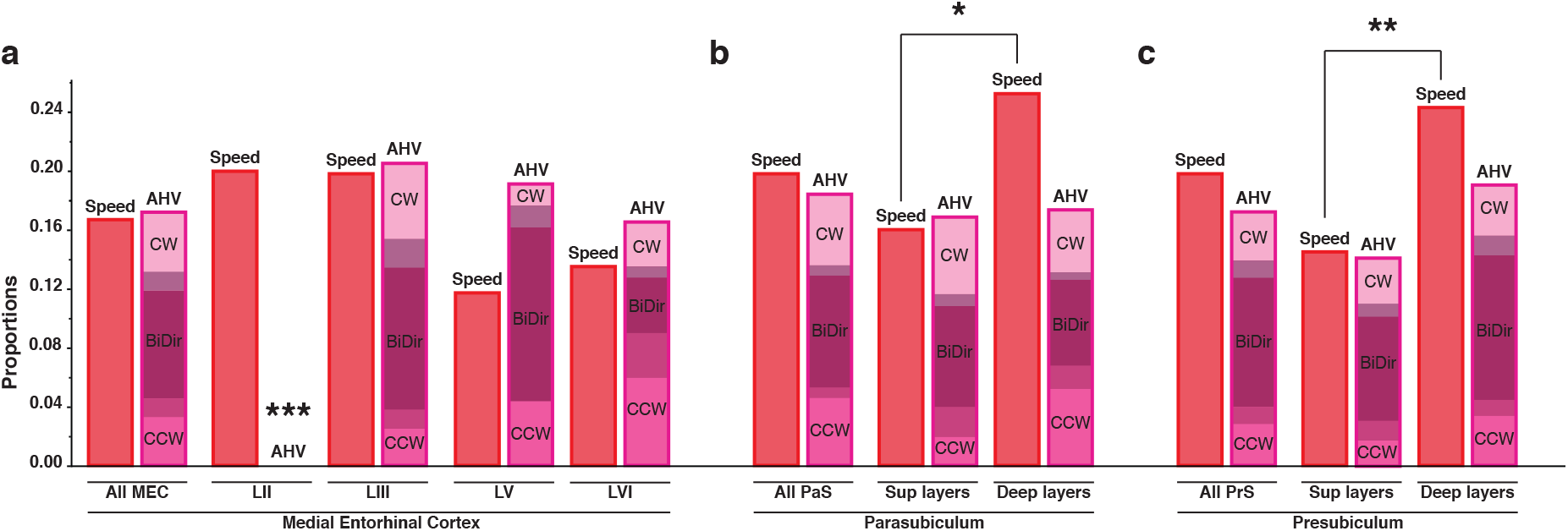
Distribution of angular head velocity and linear speed modulation across parahippocampal layer. Proportion of speed (red) and AHV cells (CW light pink, CCW dark pink, BiDir purple) across layers. Shaded areas represent overlap between CW (CCW) and BiDir cells. (**a**) Proportions in medial entorhinal cortex: MEC LII (speed: 20.0%; AHV CW: 0%; AHV CCW: 0%, AHV BiDir: 0%), MEC LIII (speed: 19.8%; AHV CW: 7.1%; AHV CCW: 3.8%, AHV BiDir: 12.8%), MEC LV (speed: 11.8%; AHV CW: 2.9%; AHV CCW: 4.4%; AHV BiDir: 13.2%) and LVI (speed: 13.5%; AHV CW: 3.7%; AHV CCW: 9%; AHV BiDir: 7.5%). Stars denote the significant absence of AHV cells in MEC LII (t-test, pvalue <0.001). (**b**) Proportions in the parasubiculum: superficial layers (speed: 16.1%; AHV CW: 6.0%; AHV CCW: 4%; AHV BiDir: 9.6%) and deep layers (speed: 25.3%; AHV CW: 4.7.9%; AHV CCW: 6.86%; AHV BiDir: 7.9%). (**c**) Proportions in the presubiculum: superficial layers (speed: 14.5%; AHV CW: 3.9%; AHV CCW: 3.1%; AHV BiDir: 9.3%) and deep layers (speed: 24.3%; AHV CW: 4.7; AHV CCW: 4.5%; AHV BiDir: 12.2%). Stars denote the significant difference in speed cells between superficial and deep layers both in PrS and PaS (t-test, *** pvalue <0.001, ** pvalue <0.01 and * pvalue <0.05 respectively).

**SF3.**
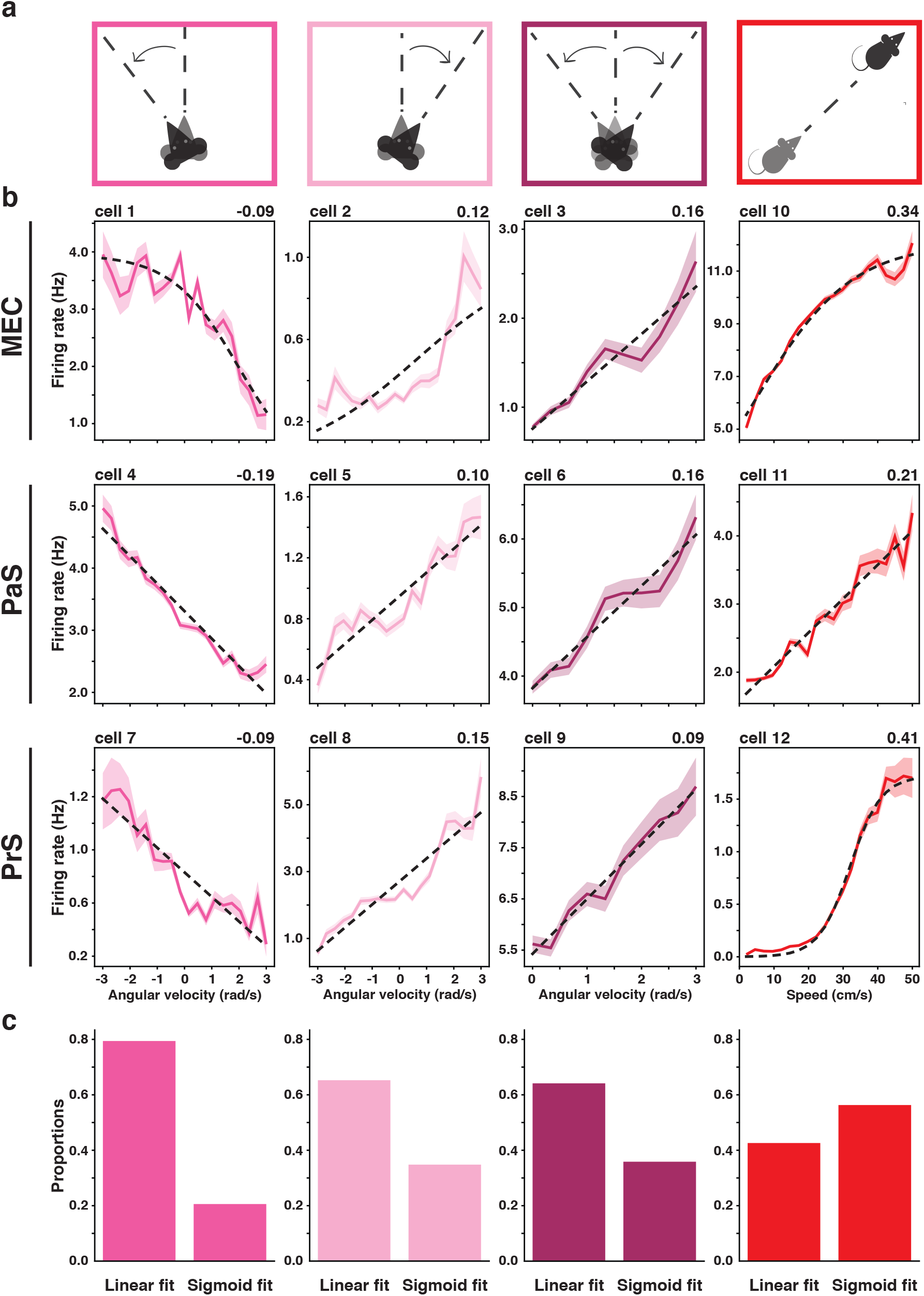
Linear and sigmoidal fit in angular head velocity (AHV) and speed modulated cells. (**a**) Schematic representation of the behavioural correlate modulating the rate. From left to right: AHV CCW (dark pink), AHV CW (light pink), AHV BiDir (purple), linear speed (red). (**b**) Examples of tuning curves of cells in MEC (top row), PaS (middle row) and PrS (bottom row), columns colour coded and arranged as in (a). Solid lines represent the average firing rate at a given value of AHV (speed) across the recording session, shaded areas represent the standard error of the mean and dotted line the best linear (sigmoidal) fit. Cell scores are reported on the top right corner and cell ID in the top left corner. (**c**) Proportion of linear and sigmoidal cell in the total population. From left to right: AHV CCW (Linear: 79.4%; Sigmoidal: 20.6%), AHV CW (Linear: 65.2%; Sigmoidal: 34.8%), AHV BiDir (Linear: 64.1%; Sigmoidal: 35.9%), speed (Linear: 42.6%; Sigmoidal: 56.3%). Note that the sigmoidal fits are often quasi-linear.

**SF4.**
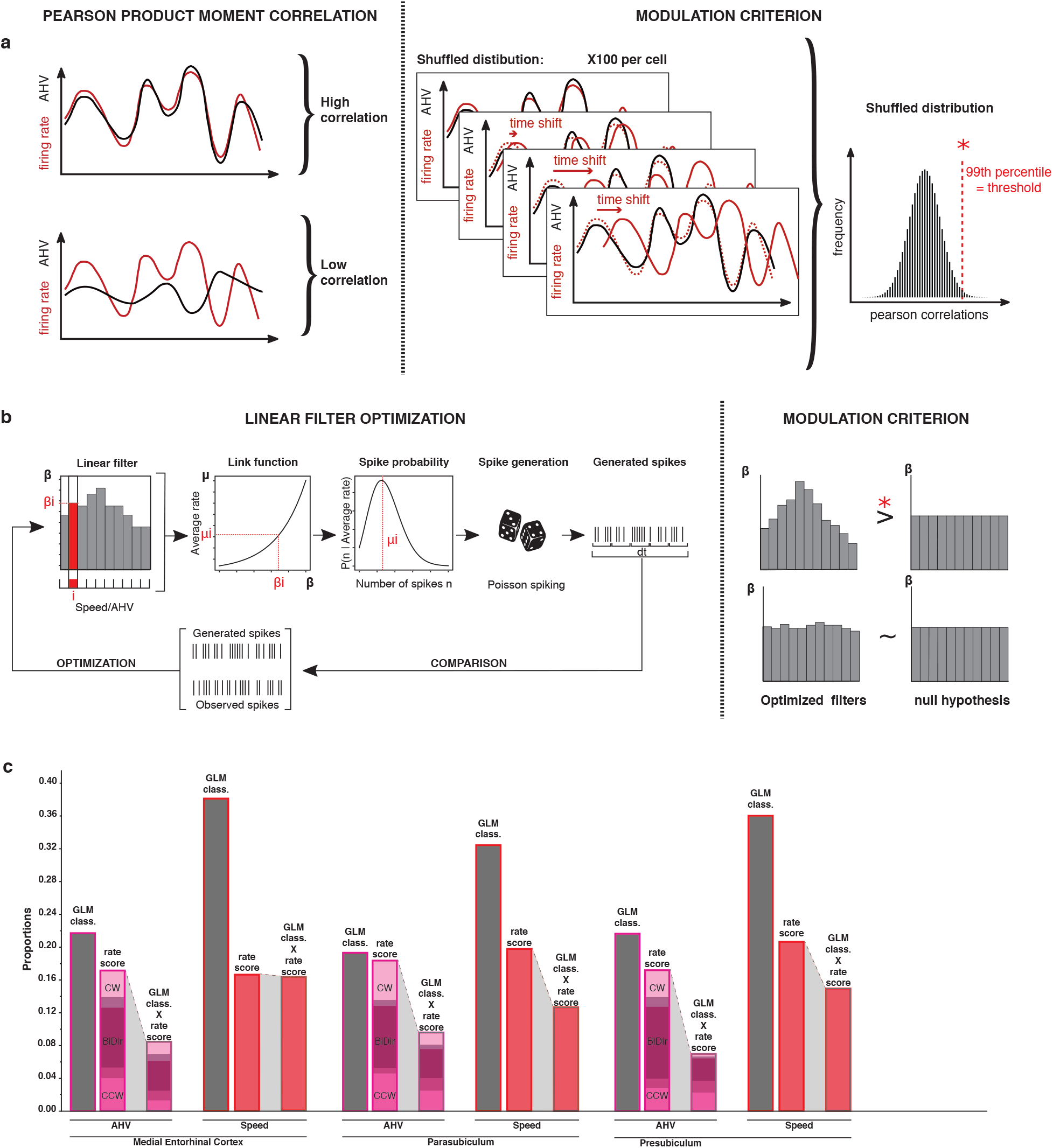
Comparison between scoring methods: correlation vs. GLM. (**a**) Schematics of the correlation method. The session-wide Pearson correlation between the firing rate of the cell (red) and the angular head velocity (black) is computed. Two examples of high (left, top) and low (left, bottom) correlations are illustrated. Significance is calculated with a shuffling procedure (right) in which the firing rate is shifted in time of a random amount, 100 times for each cell and a shuffled score is calculated. Shuffled scores of all cells in the same region are pooled together to obtain a null distribution of score. Cells are labelled as AHV modulated if their score exceed the 99th percentile of the null distribution (rightmost part). (**b**) Schematics of the generalized linear model (GLM) method. First the weights β of the GLM model are optimized with a maximum likelihood procedure (left). The value β_i_ corresponding to the bin i of the instantaneous angular head velocity determines the value µ_i_ of the instantaneous firing rate through an exponential link function. The value µ_i_ is the average of the Poisson distribution giving the probability P(n|µ_i_) to observe a certain number of spikes n in the time interval dt at which the AHV signal is sampled (20 ms in our case). This distribution is used to generate, through a Poisson spiking process, a train of spikes to be compared with the ones experimentally observed. The weights are adjusted to maximize the similarity across the whole session. The cell is labelled as modulated if the likelihood of the optimized model is significantly larger than the null model with constant β fixed at the average firing rate of the cell (right) (**c**) Intersection between correlation and GLM modulated cells in MEC (GLM modulated AHV cells: 21.7%, n=86; correlation modulated AHV cells: 16.9%, n=67; intersection: 9.6%, n=38 | GLM modulated speed cells: 38.1%, n=151; correlation modulated speed cells: 16.7%, n=66; intersection: 12.6%, n=50), parasubiculum (GLM modulated AHV cells: 19.3%, n=84; correlation modulated AHV cells: 17.2%, n=75; intersection: 6.9%, n=30| GLM modulated speed cells: 32.4%, n=141; correlation modulated speed cells: 19.8%, n=86; intersection: 14.9%, n=65), and presubiculum (GLM modulated AHV cells: 21.6%, n=131; correlation modulated AHV cells: 17.2%, n=104; intersection: 8.4%, n=51 | GLM modulated speed cells: 36%, n=218; correlation modulated speed cells: 20.7%, n=125; intersection: 16.4%, n=99). The intersection between correlation modulated and GLM modulated cells is in each case significantly larger than expected by chance (binomial test, pvalue <0.001).

<u>Comment on scoring method</u>: The correlation method and the GLM approach have rather complementary advantages and weaknesses. The former assumes that the modulation of the firing rate is linear or close to linear but does not require any further arbitrariness in the fixing of additional parameters. The latter is able to detect more general forms of modulation but is subject to choices (e.g., binning size, regularization procedure, form of the link function) that are, at least to some degree, arbitrary. Since the majority of the modulated cells are found with both methods, we proceeded in the further analysis with the simpler one, which also allows a more direct comparison with previous literature.

**SF5.**
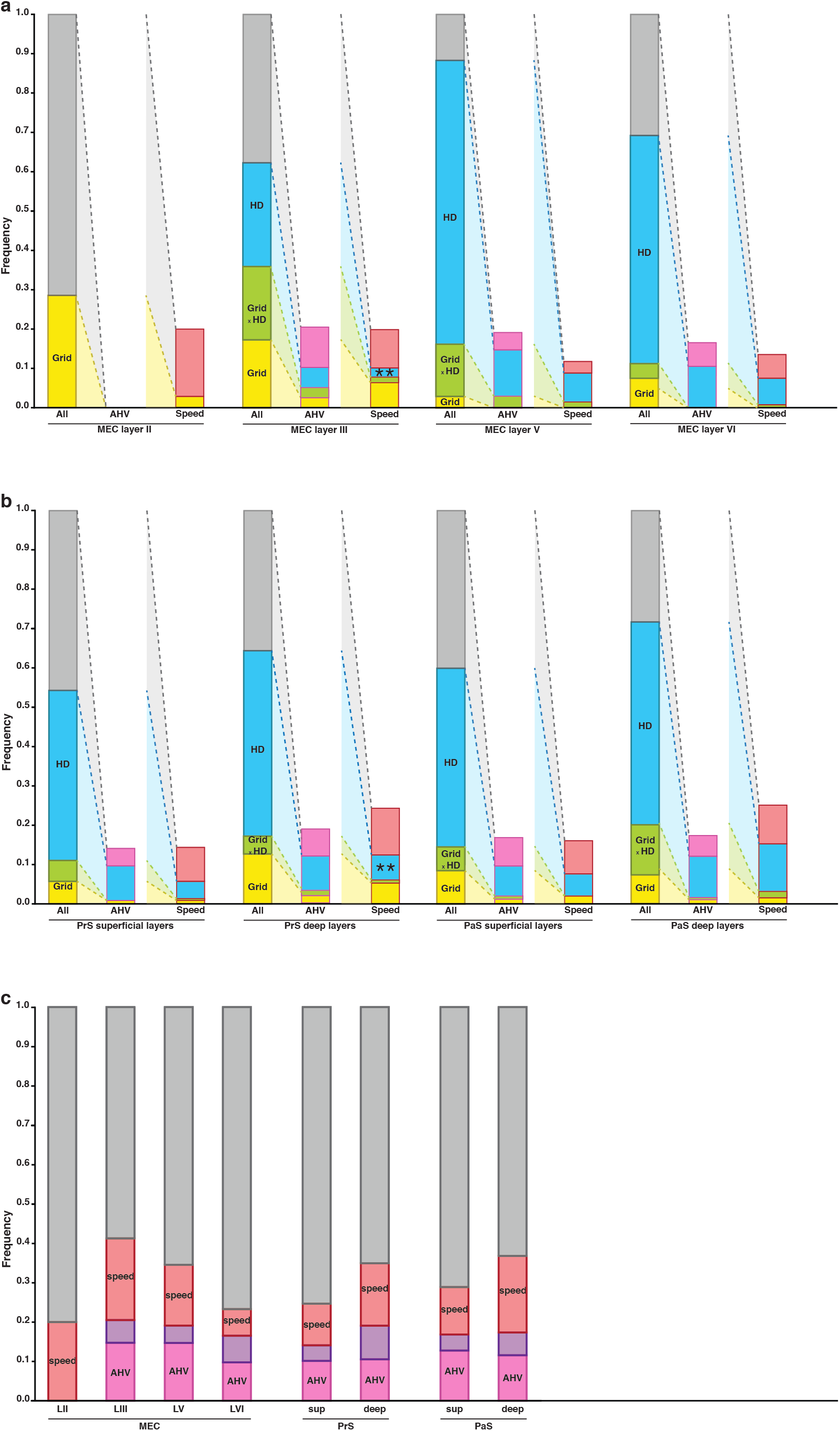
Distribution of conjunctive coding across areas and layers. (**a–b**) Proportions of grid (yellow), HD (blue), grid x HD (green) cells in the whole layer population (left bar, black outline), within the AHV cell population (central bar, pink outline) and within speed cell population (right bar, red outline). Grey bars represent cells that are neither coding for grid nor HD. Pink bars in the AHV population histograms represents AHV cells that are neither coding for grid nor HD. Red bars in the speed population histograms represents speed cells that are neither coding for grid nor HD. Stars denote a significant change in proportions of a specific type of conjunctive cells within either the AHV or the speed population from what would be expected from the layer proportions within the general population. All proportions were as expected, except for an underrepresentation of speed X HD in MEC L III and in deep Prs (t-test: pvalue <0.01). **MEC**(**a**): MEC layer II (grid: 28.6%; HD: 2%; grid x HD: 0%), layer III (grid: 35.9%; HD: 44.9%; grid x HD: 18.6%), layer V (grid: 16.2%; HD: 85.3%; grid x HD: 13.2%), and layer VI (grid: 11.3%; HD: 61.7%; grid x HD: 3.7%). **PrS**(**b**, left): PrS superficial layers (grid: 11%; HD: 48.5%; grid x HD: 5.3%), and PrS deep layers (grid: 17.2%; HD: 51.6%; grid x HD: 4.5%). **PaS**(**b**, right): PaS superficial layers grid: (14.5%; HD: 51.4%; grid x HD: 6%), PaS deep layers (grid:20%; HD: 64.2%; grid x HD: 12.6%). Note the quasi-absence of HD and AHV cells in MEC LII. (**c**) Pink and red bars here represent the whole population of AHV (pink) and speed (red) cells in a given layer. Note that those are different populations than in (a–b). Purple bars represent cells whose activity is conjunctively modulated by speed and by AHV. From left to right: MEC layer II (AHV: 0%, speed: 20%, AHV x speed: 0%), MEC layer III (AHV: 20.5%, speed: 19.9%, AHV x speed: 5.8%), MEC layer V (AHV: 19.1%, speed: 11.8%, AHV x speed: 4.4%), MEC layer VI (AHV: 16.5%, speed: 13.5%, AHV x speed: 6.8%), PrS superficial layers (AHV: 14.1%, speed: 14.5%, AHV x speed: 3.9%), PrS deep layers (AHV: 19.1%, speed: 24.3%, AHV x speed: 8.5%), PaS superficial layers (AHV: 16.9%, speed: 16.1%, AHV x speed: 4%) and PaS deep layers (AHV: 17.4%, speed: 25.3%, AHV x speed: 5.8%).

**SF6.**
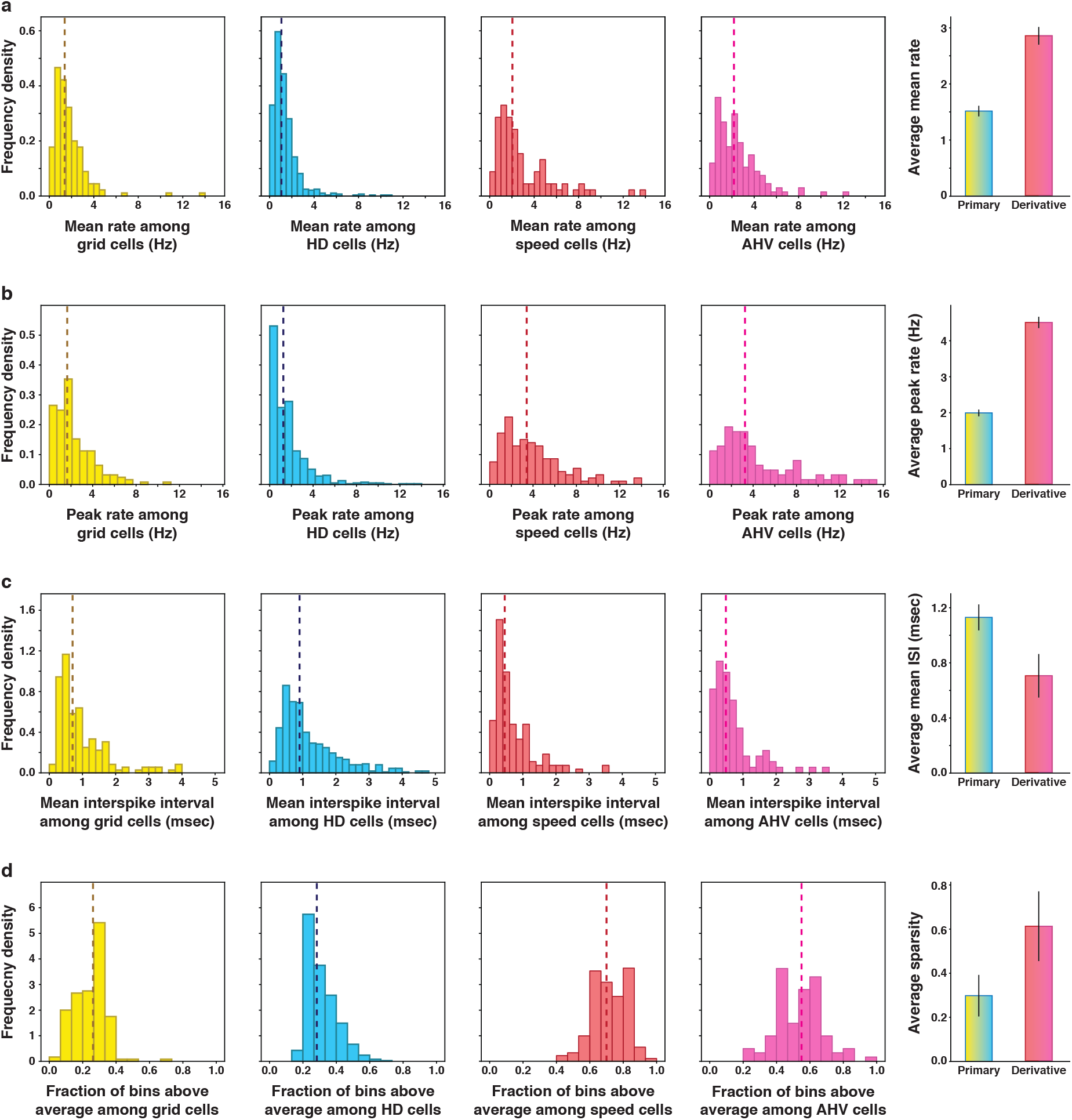
Primary and derivative correlates have different firing properties. Comparison of firing properties between cells coding for primary correlates – grid cells (yellow) and HD cells (blue) – and derivative correlates – speed cells (red) and AHV cells (pink). Vertical dashed lines indicate median values. From left to right: distribution of frequency of density of cells presenting a given range of a firing property (**a**) Distribution of the average firing rates of cells coding for motion and static signals (bin width: 0.5Hz). Median average rate for motion cells: 2.86 Hz (orange dashed line). Median average rate for static cells: 1.52 Hz (blue dashed line). (**b**) Distribution of the peak firing rates calculated as the 5th percentile of the firing rate distribution of each cell (bin width: 1Hz). Median peak rate for motion cells: 4.51 Hz (orange dashed line). Median peak rate for static cells: 1.99 Hz (blue dashed line). (**c**) Distribution of the average inter-spike interval (bin width: 0.1s). Median average inter-spike interval for motion cells: 0.71 s (orange dashed line). Median average inter-spike interval for static cells: 1.13 Hz (blue dashed line). Note that motion cells show a larger average and peak firing rate, as well as a lower average inter spike interval. This difference may be related to the nature of the coding: static cells use “place like”, sparse coding, while motion cells have a monotonic, dense response profile.

**SF7.**
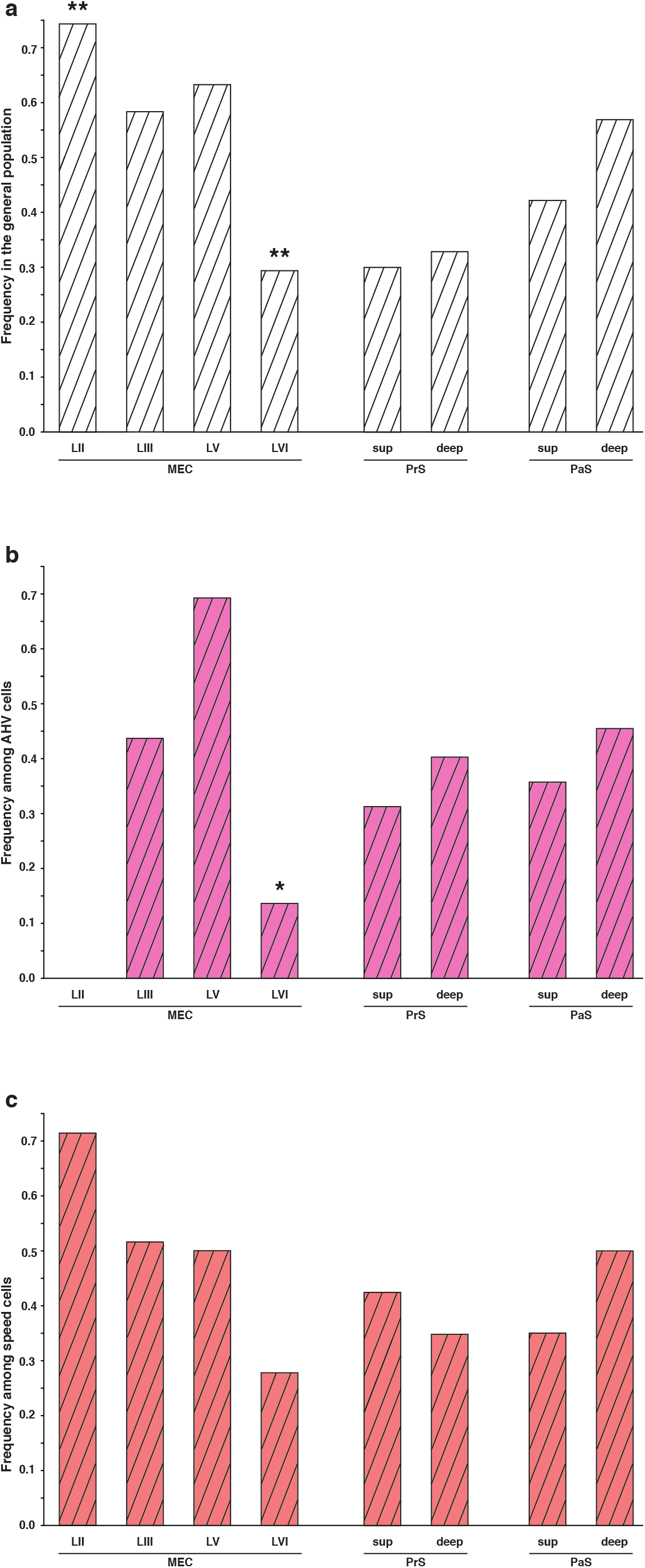
Distribution of theta modulation by layer. Dashed bars represent proportions of theta modulated cells for each category of cells considered. Cells were considered theta modulated when their mean power in a 2 Hz window centred in the peak in the 5-to 11-Hz frequency range was at least fivefold greater than the mean spectral power in the 0-to 125-Hz range. (**a**) Percentages of theta modulated cells in the whole population (white dashed bars). From left to right: MEC LII (74.3%), MEC LIII (58.3%), MEC LV (63.2%), MEC LVI (29.3%), PrS superficial layers (29.9%), PrS deep layers (32.8%), PaS superficial layers (42.2%) and PaS deep layers (56.8%). Stars denote a significant difference in proportion of theta modulated cell in the layer considered, compared to the average theta modulation across all layers (t-test, ** pvalue <0.01). (**b**) Percentages of theta modulated cells in the AHV population (pink dashed bars). From left to right: MEC LII (0%), MEC LIII (43.8%), MEC LV (69.2%), MEC LVI (13.6%), PrS superficial layers (31.3%), PrS deep layers (40.2%), PaS superficial layers (35.7%) and PaS deep layers (45.5%). Stars denote a significant difference in proportion of AHV theta modulated cell from what would be expected given the average theta modulation in that specific layer (t-test, * pvalue <0.05). (**c**) Percentages of theta modulated cells in the speed population (red dashed bars). From left to right: MEC LII (71.4%), MEC LIII (51.6%), MEC LV (50%), MEC LVI (27.8%), PrS superficial layers (42.4%), PaS deep layers (34.7%), PaS superficial layers (35%) and PaS deep layers (50%).

## Methods

### Subject and surgeries

All the data presented here have been previously published^30^ but was re-analysed for this manuscript. The neuronal activity was recorded from 28 male Long-Evans rats (3–5 months old, 350–450 g at implantation, housed and food deprived as in described previously). All experiments were approved by the National Animal Research Authority of Norway. Tetrodes configuration and surgical implantations are described in previously published work^30^.

### Data acquisition and training procedures

General data acquisition procedures have been described previously^30^. In brief, rats were trained to collect food crumbs thrown randomly into a 50-cm-high square or circular box with black floor and black walls surrounded by black curtains. Each trial lasted 10 mins.

### Spike sorting

The spike detection in the local field potential and sorting was performed as previously described^30^. Spike sorting was performed offline using graphical cluster-cutting software.

### Estimation of the behavioural correlates

#### Position

The position of the animal was estimated from the coordinates of two light-emitting diodes (LEDs) on the head of the animal. The X and Y coordinates of both the LEDs where smoothed with a gaussian filter with a 250 ms standard deviation, chosen to match the smoothing performed on the firing rate (see below), and the average between the two LED positions was used as the position of the animal.

#### Head Direction

The head direction (HD) was calculated as the angle between the line connecting the small LED to the big LED and the x axis. HD is expressed in radians, 0 meaning that the rat head is aligned with the x-axis, facing right.

#### Linear speed and angular head velocity

Speed was calculated as the modulus of the vector difference between the position at time t and the position at time_t+1_. Angular head velocity was calculated as the signed difference between the head direction at time t and the head direction at time_t+1_. The absolute value of the angular head velocity was used for the scoring of bidirectional angular head velocity cells (see below). No further smoothing was applied.

### Firing rate calculation

Instantaneous firing rate was obtained dividing the whole session in bins of 20 ms, coinciding with the frames of the tracking cameras. The spike count in each time bin was then calculated and divided by the temporal width to obtain the rate. The rate profile was smoothed with a 250 ms wide Gaussian filter.

### Speed filtering

The analysis on speed and angular velocity was performed on movement periods, defined as the ones in which the animal speed was >2 cm/s. A speed filter was applied on the timeseries of each correlate, discarding the time points for which the instantaneous speed was below 2 cm/s, that were excluded in the subsequent analysis.

### Rate maps and tuning curves

#### Spatial rate maps

The histograms for spike count and time spent in each location were constructed using equally spaced bins of 2-cm linear size. Each bin of the rate map was obtained as the ratio between spike count and time spent, smoothed with a gaussian filter with standard deviation of 4 cm.

#### Directional rate maps

The histograms for spike count and time spent facing each direction were constructed using equally spaced bins of size 6 degrees. Each bin of the rate map was obtained as the ratio between spike count and time spent, smoothed with a gaussian filter with standard deviation of 6 degrees.

#### Speed and angular velocity tuning curves

For tuning curve construction, the correlate was divided in equally spaced bins. For speed, 20 bins spanned the range between 2- and 50-cm/s (bin width: 2.4 cm/s), for angular velocity the range −3-, +3-rad/s was again divided into 20 bins (bin width: 0.15 rad/s). The firing rate in each bin was calculated as the average of the instantaneous firing rate values falling in the each given bin. A gaussian smoothing window with standard deviation 0.15 rad/s for angular velocity and 2.4 cm/s for speed was applied.

### Shuffling

Chance-level statistics was calculated for a given variable W through a shuffling procedure. For each repetition, the firing rate time series was time shifted of a random interval of at least 30 seconds, with the end of the trial wrapped to the beginning.

This procedure was repeated 100 times for each cell, and the shuffled score for variable W was calculated for each instance to compose the chance level statistics.

For cell classification, all shuffled data from the same region were pooled together and the 99th percentile of the distribution was used as a classification threshold.

### Measure used for cell type classification

#### Speed Score

The speed score was defined as the Pearson product-moment correlation between the cell’s instantaneous firing rate and the instantaneous speed of the animal, across the whole recording session. This yields a score ranging from −1 to +1.

#### Unidirectional angular velocity score

The unidirectional angular velocity score was defined as the Pearson product-moment correlation between the cell’s instantaneous firing rate and the instantaneous angular velocity of the animal. Positive values of angular velocity correspond to clockwise head movement. Cells that had a score greater than the 99th percentile of the shuffled distribution were classified as clockwise modulated (CW), while cells whose score was lower than the 1st percentile were classified as counterclockwise modulated (CCW): they significantly code for head movement in the counterclockwise direction. CW and CCW populations are mutually exclusive by construction.

#### Bidirectional angular velocity score

The bidirectional angular velocity score was defined as the Pearson product-moment correlation between the cell’s instantaneous firing rate and the absolute value of the instantaneous angular velocity of the animal. Cells in this population increase their firing rate in response to head movement regardless of the direction. The unidirectional and bidirectional angular velocity scores are not mutually exclusive by construction. A cell with a strong ramping up of the activity for positive value of AHV, and a mild sensitivity to negative values, for example, could be selected as both a bidirectional and a unidirectional CW cell. A strong modulation from both negative and positive angular velocities would be picked up only by the bidirectional score, while a strictly monotonic increase (or decrease) along the whole range of velocities would give a high unidirectional score.

#### Mean vector length (head-direction score)

The mean vector length score is calculated from the head-direction tuning map of a given cell as the sum:

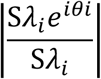

where *θ_i_* is the orientation in radians associated with bin *i* and *λ_i_* is the firing rate in the bin. The sums run over all N directional bins, and the modulus of the resulting complex number is taken. Head direction was binned in bins of 6 degrees and smoothed with a gaussian filter with a standard deviation of 6 degrees.

#### Grid score

The grid score was calculated from the spatial autocorrelogram of a given cell with a procedure similar to ^66^. After exclusion of the centre of the autocorrelogram, the Pearson correlation of the autocorrelogram rotated by 30,60,90, 120 and 15 degrees (+-3 degrees offsets) was considered. Only bins closer to the centre than an outer radius *s* were included in the calculation of the correlation. Given *s*, the grid score was defined as the difference between the average of the maximum correlations around 60 and 120 degrees (+-3 degrees offsets) and the average of the minimum correlations around 30,90 and 150 degrees (+-3 degrees offsets). The final grid score of the cell was then defined as the maximum grid score over values of s ranging from twenty to forty bins, computed at intervals of one bin.

#### Theta index

Theta modulation of individual cells was estimated from the frequency power spectrum of the spike-train autocorrelation histogram of the cell.

A cell was defined to be theta modulated if the mean power in a 2 Hz window centred in the peak in the 5-to 11-Hz frequency range was at least fivefold greater than the mean spectral power in the 0-to 125-Hz range.

### Estimation of the significance of overlaps between cell populations

The observed overlaps between cell populations were compared to the ones that would result from the statistical null hypothesis of independent random assignment with a two-sided binomial test. The probability of observing an overlap of size k between two populations of sizes *N_a_* and *N_b_*, independently drawn from a total number of cell N is given by:

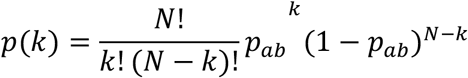

Where *p*_*ab*_ = *p*_*a*_x *p*_*b*_ and *p*_*a*_ = *N*_*a*_/*N*, *p*_*b*_ = *N*_*b*_/*N*.

### Information analysis

The information per spike conveyed by each cell about the correlate of interest (speed or angular head velocity) was calculated using the formula:

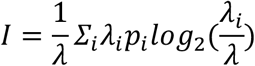

Where *i* is the index of the correlate bin, *p*_*i*_ is the probability of observing the correlate in bin *i* (i.e. the normalized occupancy), *λ*_*i*_ is the average firing rate of the cell in bin *i*, and *λ* is the average firing rate of the cell.

Speed was divided in 2 cm/s bins in the range 2-50 cm/s (as in all analysis, stillness periods were excluded), while angular head velocity was divided in 0.5 rad/s bins, in the range −5-5 rad/s.

Cells were considered to carry significant information about the correlate if the observed information rate exceeded the 99th percentile threshold of the null distribution obtained by shuffling the cell firing rate values (1000 shuffles per cell).

### Generalised linear model (GLM) analysis

We analysed the effect of each correlate (speed and angular head velocity) with a linear-nonlinear Poisson spiking GLM model.

This model assumes that the firing rate of the cell depends on the value of the correlate as:

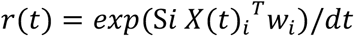

Where *X*_*i*_(*t*) is a one-hot vector (i.e., a vector with only one non-null element) indexing which value the correlate is taking at time*t*, and *w*_*i*_ are the coefficients of a linear filter quantifying the contribution of each value of the correlate to the firing rate of the cells.

The model is fitted using the python module statsmodel.api, which finds the set of parameters *w*_*i*_ maximising the log-likelihood of the observed spikes, subject to an elastic-net regularization constraint.

To perform the fitting procedure, the speed values have been binned in 10 bins in the range 2-50 cm/s, and the angular velocity values divided in 10 bins in the range −3-3 rad/s.

Cells were considered significantly modulated by a correlate if the log-likelihood of the best fit was significantly larger than the value obtained with only the average firing rate as a predictor. Significance was estimated with a 10–fold bootstrapping procedure to extract the confidence interval of the observed log-likelihood.

### Tuning curve fitting

Two different functional forms were fitted and compared to the tuning curve of modulated cells.

A linear model:

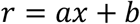

And a sigmoid model:

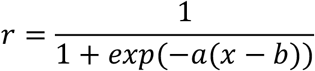

Here x is the value of the correlate (speed or angular head velocity) and r is the average firing rate of the cell at that value of x. Tuning curves were rescaled by their maximum value, in order to match the two model by number of parameters.

The R2 fitting scores were then compared for each cell. Cells with a linear R2 greater than the sigmoid R2 were classified as linear, and vice versa.

To quantify the number of cells that showed modular coding (i.e., an increased rate in a particular speed or angular velocity band), we fitted their rescaled tuning curves with a gaussian profile:

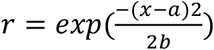

Cells were labelled as gaussian if the fit yielded a R2 score > 0.5 and the average a of the gaussian laid within the tuning curve interval (2-50 cm/s for speed, −3-3 rad/s for AHV).

### Sparsity calculation

To quantify the sparsity of the firing of primary and derivative cells, we calculated their tuning curves (as described above) as a function of speed (for speed cells), of AHV (for AHV cells), head direction (for HD cells) and position (for grid cells).

We then calculated as the sparsity the percentage of the bins of the tuning curve that had a firing rate value larger than the average overall firing rate of the cell.

